# Evaluation of Potentially Toxic Elements in Soil and Potential Ecological Risks Generated by Environmental Liabilities in Tacna, Peru

**DOI:** 10.1101/2024.09.21.614289

**Authors:** César Julio Cáceda, Gisela Maraza, Gabriela de Lourdes Fora, Diana Galeska Farfan, Edwin Obando, Fulvia Chiampo, Milena Carpio

## Abstract

Environmental liabilities continue to pose an unresolved concern for administrators due to their high potential for ecosystem contamination. This research focuses on assessing the content of potentially toxic elements, the degree of contamination, and potential ecological risks in abandoned mining areas that formerly exploited sulfur and copper. The results showed elevated concentrations of Arsenic (1,102 mg/Kg), Cadmium (271 mg/Kg), Lead (15,961 mg/Kg). The presence of flora, fauna activity, rivers, and rural communities near the mining sites was observed, indicating significant environmental risks. The evaluated mining environmental liabilities (MELs) lack direct responsible parties, thus requiring the Peruvian government to assume remediation responsibilities. To date, no mitigation actions have been taken, primarily due to the absence of a situational diagnosis. Concerning contamination indices, such as the Geoaccumulation Index, Contamination Degree, Pollution Load Index, Contamination Load Coefficient, and Potential Ecological Risk Index, all areas exhibited some form of contamination, indicating high environmental risks. A preliminary risk assessment associated with the presence of mining environmental liabilities has been conducted, marking this research as the first of its kind in the southern region of Peru. This assessment provides administrators with crucial information to establish priorities for implementing remediation plans aimed at reducing pollutant loads. The findings underscore the urgent need for comprehensive contamination assessments and the development of effective management practices, including the implementation of a monitoring program to safeguard soils affected by mining activities. Additionally, it is essential to design various technological strategies to restore degraded ecosystems, thereby protecting rivers, agricultural zones, and nearby rural communities.

## Introduction

Mining is one of the oldest and most significant anthropogenic activities globally, remaining a crucial economic sector for numerous countries (1,2). However, inadequate waste management during mining activities has resulted in negative effects due to the accumulation of hazardous waste (3), leading to environmental contamination affecting

Mining is one of the oldest and most significant anthropogenic activities globally, remaining a crucial economic sector for numerous countries (1,2). However, inadequate waste management during mining activities has resulted in negative effects due to the accumulation of hazardous waste (3), leading to environmental contamination affecting various ecosystems, primarily by heavy metals and metalloids (4,5). This situation has caused social conflicts and health problems (1,6). Historically, areas used for mining received no remediation, mainly due to the lack of environmental regulations. Few countries had mining regulations and government recovery programs, resulting in the accumulation of Mining Environmental Liabilities (MELs) worldwide (7,8).

It is estimated that there are over a million abandoned mines globally, including galleries, alluvial mines (7), tailings, topography, and unused wells (9), generating toxic waste (7,10,11). In Latin America, more than 1,266 Mining Environmental Liabilities (MELs) have been officially reported in the Plurinational State of Bolivia, 492 in Chile, 522 in Colombia, and 7,956 in Peru (12,13). Most of these environmental liabilities lack an identified responsible party (14), leading to social conflicts (15). Peru faces a significant challenge regarding MELs. Despite being the first country in Latin America to establish a legal framework for MEL sites in 1993 (16), a significant number of these MELs originated before regulation, when standards were less stringent than they are today (8,17). These Mining Environmental Liabilities are defined as facilities, waste, or residues generated by mining operations that are currently abandoned or inactive. They include effluents, emissions, and residue remnants from mining activities, posing long-term harm to the population and the developed ecosystem (8,18).

Environmental pollution caused by potentially toxic elements (PTE) presents a significant challenge due to their high toxicity, long persistence, and lack of biodegradability (19–21). These contaminants tend to disperse into surrounding areas, including agricultural fields, vegetation, and forest landscapes (22), causing soil nutrient loss through erosion and toxic gas emissions (9,23–26). Additionally, they can alter the biological and physicochemical properties of the soil, inducing harmful effects on various plant physiological processes (27,28), and their accumulation leads to degradation and the risk of contamination of surface (29), groundwater (30), and ecosystem (31), eventually entering the food chain and becoming detrimental to human health (28,32–35). Exposure through water, soil, and air can result in acute or chronic intoxications, and the bioaccumulation of these potentially toxic elements can trigger adverse effects in various tissues and organs of the body (36). Moreover, most of these compounds possess carcinogenic properties (37), and according to the International Agency for Research on Cancer (37), Arsenic (As), Cadmium (Cd), Chromium (Cr), and Nickel (Ni) are Category 1 heavy metals regarding this pathology. In Russia, high concentrations of Zn, Ni, Pb, Cu, and Cd increase due to the interaction of waste rocks, atmospheric precipitation, and mine acid drainage (38,39). In Spain and China, the dispersion of these elements has contaminated surrounding areas, agricultural fields, urban soils, and forest landscapes (23–26). In Ecuador, tailings deposits are a concern for the population and the ecosystem due to the potential risk of heavy metal contamination towards rivers, which are a water source for nearby communities (29). In Peru, high concentrations of toxic elements Pb, As, Zn, and Cd, resulting from leaching, reach watersheds, affecting biodiversity (1).

In various countries, these events generate constant environmental concerns as these potentially toxic elements (PTE) associated with environmental factors lead to the mobilization of metals and waste (40–42), posing an ecological risk. Therefore, identifying hazards and characterizing risk (43) are essential for assessing the extent of soil contamination (44). In this regard, various indices are employed to assess the presence and concentration of anthropogenic contaminants in the environment, such as the geoaccumulation index (*I-geo*), pollution factor (C*_f_*), contamination index (PI) (45), pollutant load index (PLI), contamination degree (C*deg*) (46), and ecological risk factor (RI) (47). Given the intricacies of the landscape and surroundings, it becomes imperative to integrate multiple assessment methods to determine the contamination levels caused by potentially toxic elements.

The aim of this study is to evaluate the degree of contamination from toxic compounds generated by mining environmental liabilities (MELs) and analyze potential environmental risks. The focus lies on the concentration of toxic compounds and risk assessment utilizing indices such as the Geoaccumulation Index, Pollution Factor, Pollutant Load Index, Contamination Degree, Ecological Risk Factor, and Potential Ecological Risk Index. This will facilitate the analysis and communication of environmental information, leading to the development of mitigation strategies for soils and waters contaminated with potentially toxic elements, as well as the containment of the potential hazards they pose.

## Materials and methods

### Study Area

The study area corresponds to Environmental Mining Liabilities (MELs) located in the districts of Palca, Pachía, Candarave, and Susapaya in the Tacna Region (Fig 1). Currently, approximately 165 MELs are registered, with the main subtypes identified as mine entrances, mine waste, tailings, and infrastructure (13). These liabilities were generated by mines engaged in the extraction of copper and sulfur, resulting in a significant contamination issue, given that many of them are situated in close proximity to water bodies, near populated centers, and rural communities. Due to the mountainous physiography, there are areas with average slopes ranging between 35% and 50%. Additionally, the highly variable climatic conditions lead to the intermittent activation of rivers near the environmental liabilities, posing a risk to the surrounding ecosystem.

**Fig 1.**
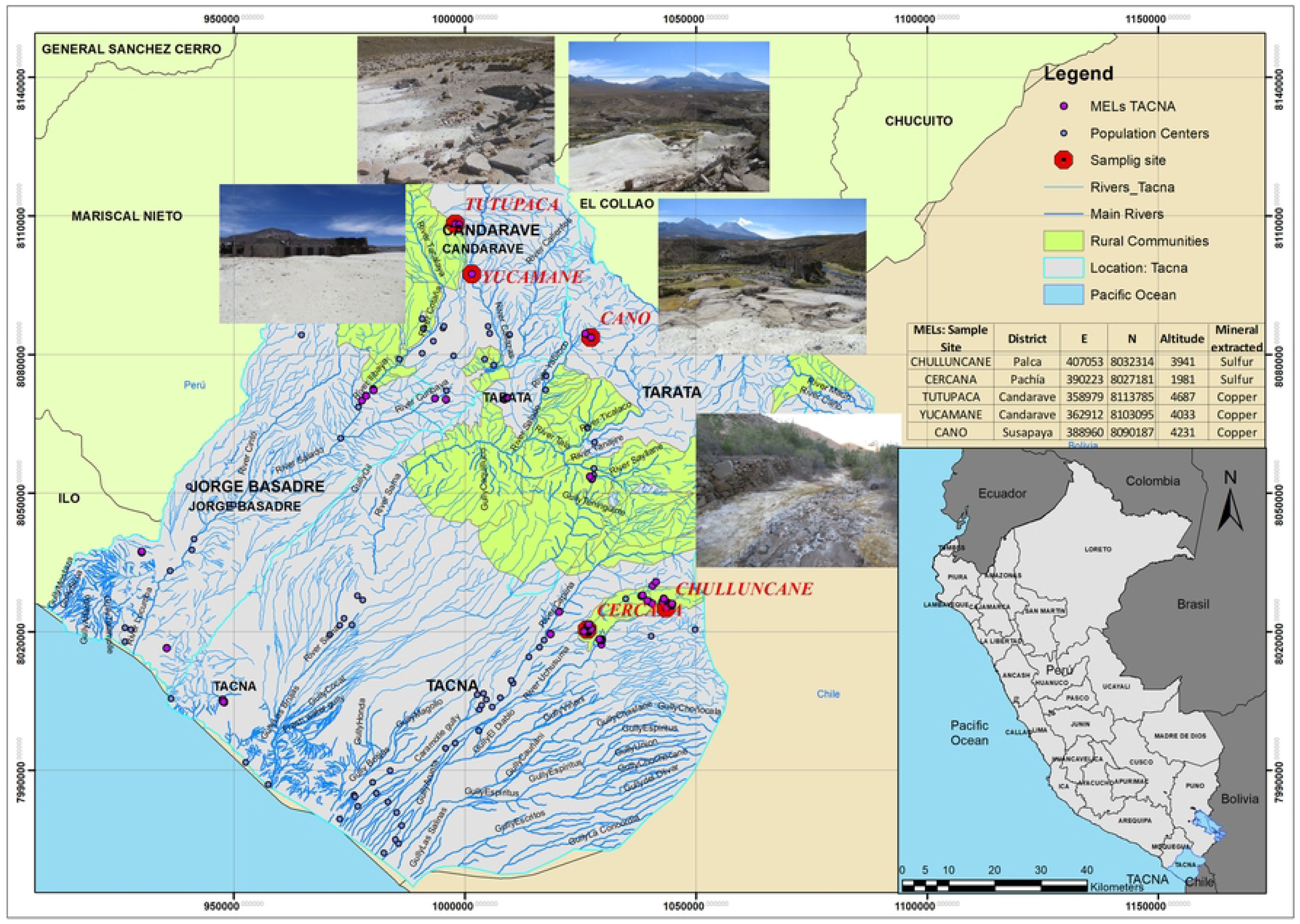
Location of environmental liabilities material in the Tacna Region, Peru. MELs: Mining environmental liabilities, Sample site: Tutupaca, Yucamane, Cano, Chulluncane, Cercana.

Furthermore, the following criteria were employed to depict the actual situation of the MELs: the name of the environmental liability, geolocation, mining activity, type of waste, distance to water bodies (within 0 to 100 meters), presence of vegetation, signs of wildlife, distance to the affected site (within 0 to 100 meters) (48–50), populated centers, and rural communities.

### Sample Collection

In each study area, three sampling points were randomly selected for each environmental liability (19), at a depth of 0-10 cm (51). Approximately 500 g of environmental liability material was collected, sealed, and stored in dense polyethylene bags (52). Subsequently, the samples were transported to the laboratory for further analysis of inorganic components in accordance with the environmental quality standards for industrial soils (18). Additionally, soil samples without anthropogenic intervention (referred to as "native soil") were collected (53), resulting in a total of 20 samples. The GPS coordinates of the sampling sites are presented in Fig 1.

### Concentration of Metals and Free Cyanide in Mining Environmental Liabilities

The concentration of the following elements was determined in a laboratory accredited by the National Quality Institute (INACAL). The analyzed metals included arsenic, barium, cadmium, lead, chromium, and mercury, utilizing the mass spectrometry method (54) with detection limits of 0.02 mg/kg, 0.04 mg/kg, 0.02 mg/kg, 0.02 mg/kg, 0.2 mg/kg, and 0.03 mg/kg, respectively. Free cyanide was also assessed (55) with a detection limit of 0.05 mg/kg, and hexavalent chromium was measured using the ion chromatography method (56) with a detection limit of 0.04 mg/kg. Additionally, the pH of the samples was determined according to the US EPA 9045D (57) using a pH meter (Hanna Instruments HI3222) and electrode (Hanna Instruments HI1043-B), with a detection limit of 0.001 pH.

### Evaluation of Potential Environmental Risks

For the assessment of potential ecological risks and pollution levels, the following indices were determined: Geoaccumulation Index (*I-geo*) (57), Pollution Load Index (28,46), Pollution Index, Contamination Degree (58), Modified Contamination Degree, Ecological Risk Factor (*Eri*), and Potential Ecological Risk Index (46). These indices are crucial and widely employed to evaluate contamination levels in soil and sediments, serving as valuable tools in establishing potential ecological risks (59–63).

#### Geoaccumulation Index (I-geo) for Assessing Soil Metal Contamination

The Geoaccumulation Index (*I-geo*) was employed to evaluate the contamination levels of each metal on the soil surface. This index takes into account both the background value and analyzes the degree of contamination. The *I-geo* was calculated as described by Mueller (58):

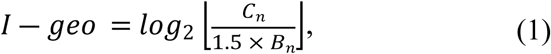

Where *I – geo* is Geoaccumulation Index, *C*_*n*_ is Concentration of the examined metal in the soil sample, *B*_*n*_ is Background value, and 1.5 as a correction factor to account for potential variations in specific metal background values.

According to Mueller (1986), *I-geo* values are classified into seven groups: For *I – geo* ≤ 0, Non-contaminated; for 0 ≤ *I – geo* ≤ 1, Non-contaminated to moderately contaminated; for 1 ≤ *I – geo ≤ 2,* Moderately contaminated; for 2 <*I-geo* ≤3, Moderately to heavily contaminated; for 3 ≤ *I – geo* ≤ 4, Highly contaminated; for 4 ≤ *I – geo* ≤ 5, Heavily to extremely contaminated; and for 5 < *I – geo*, Extremely high contamination levels.

#### Pollution Factor

The Pollution Factor (*C_f_*) was employed to quantify the contamination from hazardous compounds (46,64). It serves as an effective tool for monitoring pollution over a period of time (58). The *C_f_*value for each element was calculated using the following equation:

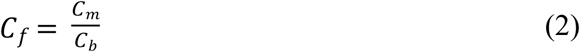

Where *C_f_:* is Pollution Factor; *C_m_*is Concentration of the element in the collected soil samples, *Cb* is Background value of the specific metal (native soil, Table 2). According to Buat-Menard and Chesselet (1979), when *C_f_* < 1, Low pollution factor; 1 ≤ *C_f_* < 3, Moderate pollution factor; 3 ≤ *C_f_* < 6, Considerable pollution factor, and *C_f_* ≥ 6, Very high pollution factor (65).

#### Degree of Contamination

The degree of contamination (*Cdeg*) is the sum of the pollution factors for the investigated elements, and it was calculated using the following equation (46).

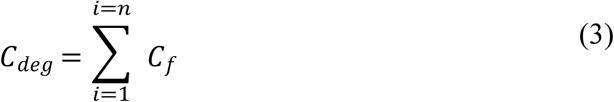

Where the following classification was proposed: *Cdeg* < 6, indicates a low degree of contamination; 6 < *Cdeg* < 12, represents a moderate degree of contamination; 12 < *Cdeg* < 24, signifies a considerable degree of contamination; and *Cdeg* > 24, indicates a high degree of contamination, suggesting severe anthropogenic pollution.

#### Modified Contamination Degree (*mCdeg*)

The modified contamination degree was introduced to estimate the overall contamination level at a site (66). The *mCdeg* was calculated using the following equation:

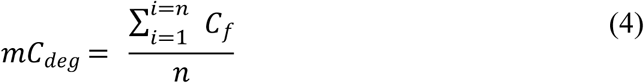

Where *n* is number of elements analyzed, *mCdeg* < 1.5, indicates a null to very low degree of contamination; 1.5 < *mCdeg* < 2, represents a low degree of contamination; 2 < *mCdeg* < 4, signifies a moderate degree of contamination; 4 < *mCdeg* < 8, indicates a high degree of contamination; 8 < *mCdeg* < 16, reflects a very high degree of contamination; 16 < *mCdeg* < 32, suggests an extremely high degree of contamination; *mCdeg* ≥ 32, indicates an ultra-high degree of contamination.

#### Pollution Load Index

The Pollution Load Index (*PLI*) integrates multiple pollutant factors to determine the contamination potential, representing the relationship between the production level and the pollutant load of each element concerning the total number of parameters analyzed (67,68). The pollution load index was calculated from the values of the pollution factor of individual elements using the following equation.

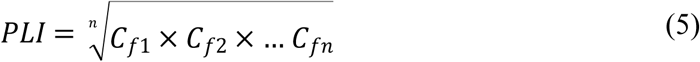

Where *PLI* is Pollution Load Index, *n* is number of metals evaluated, *C*_*fn*_is Pollution factor of a specific metal. *PLI* < 1, Uncontaminated; *PLI* =1, Slightly contaminated; *PLI* >1, Environment deterioration.

#### Assessment of Ecological Risk

The potential ecological risk index was proposed by Hakanson (1980) to evaluate the characteristics and environmental behavior of heavy metal contaminants in sediments (46).

##### Ecological Risk Factor (*Eri)*

The primary function of this index is to indicate the contaminating agents and prioritize pollution studies. The ecological risk factor (*Eri*) is calculated using the following equation. *C_f_* is the pollution factor, and *T_ri_* is the toxic response factor, representing the potential danger of heavy metal pollution by indicating the toxicity of specific heavy metals and the environmental sensitivity to pollution. According to the standardized toxic response factor (*T_ri_*) proposed by Hakanson (1980), Hg, Cd, As, Co, Cu, Pb, Cr, and Zn have toxic response factors of 40, 30, 10, 5, 5, 5, 2, 1, respectively

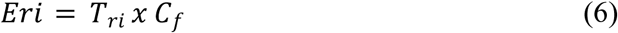

Where risk factor is *Eri* < 40, Low ecological risk; 40 < *Eri* < 80, Moderate ecological risk; 80 < *Eri* < 160, Considerable ecological risk; 160 < *Eri* < 320, High ecological risk; and 320 < *Eri*, Very high ecological risk.

##### Potential Ecological Risk Index (*IR*)

The potential ecological risk index for various heavy metals in the soil was analogous to the contamination level, defined as the sum of a singular potential ecological risk factor. This method comprehensively considers the synergy, toxic level, concentration of heavy metals, and the ecological sensitivity of the heavy metals (47). The potential ecological risk index for all measured heavy metals was calculated using the following equation.

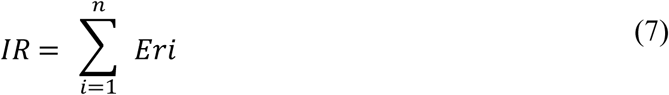

Where for the potential ecological risk index, the following terminology was utilized is *IR* < 150, Low ecological risk or low ecological contamination level; 150 ≤ *IR* < 300, Moderate ecological contamination level or moderate ecological risk; 300 ≤ *IR* < 600, Considerable ecological risk or serious ecological contamination level; and *IR* > 600, Very high ecological risk or severe ecological contamination level (46,69,70).

## Results

### Characterization of the Area and Environmental Liabilities

The results of the area characterization describe the sampling site, vegetation, peasant communities, and bodies of water detailed in Table 1. The presence of shrub, bush, and grassland flora near mining exploitation areas was reported, along with evidence of rodent activity, traces left by camelids and wild cats, pens for domestic animal breeding, and inhabitants engaged in pastoral activities. Additionally, in the Environmental Management Plans (EMPs), water infiltrations were observed in areas where tailings and waste rock are located.

**Table 1.**
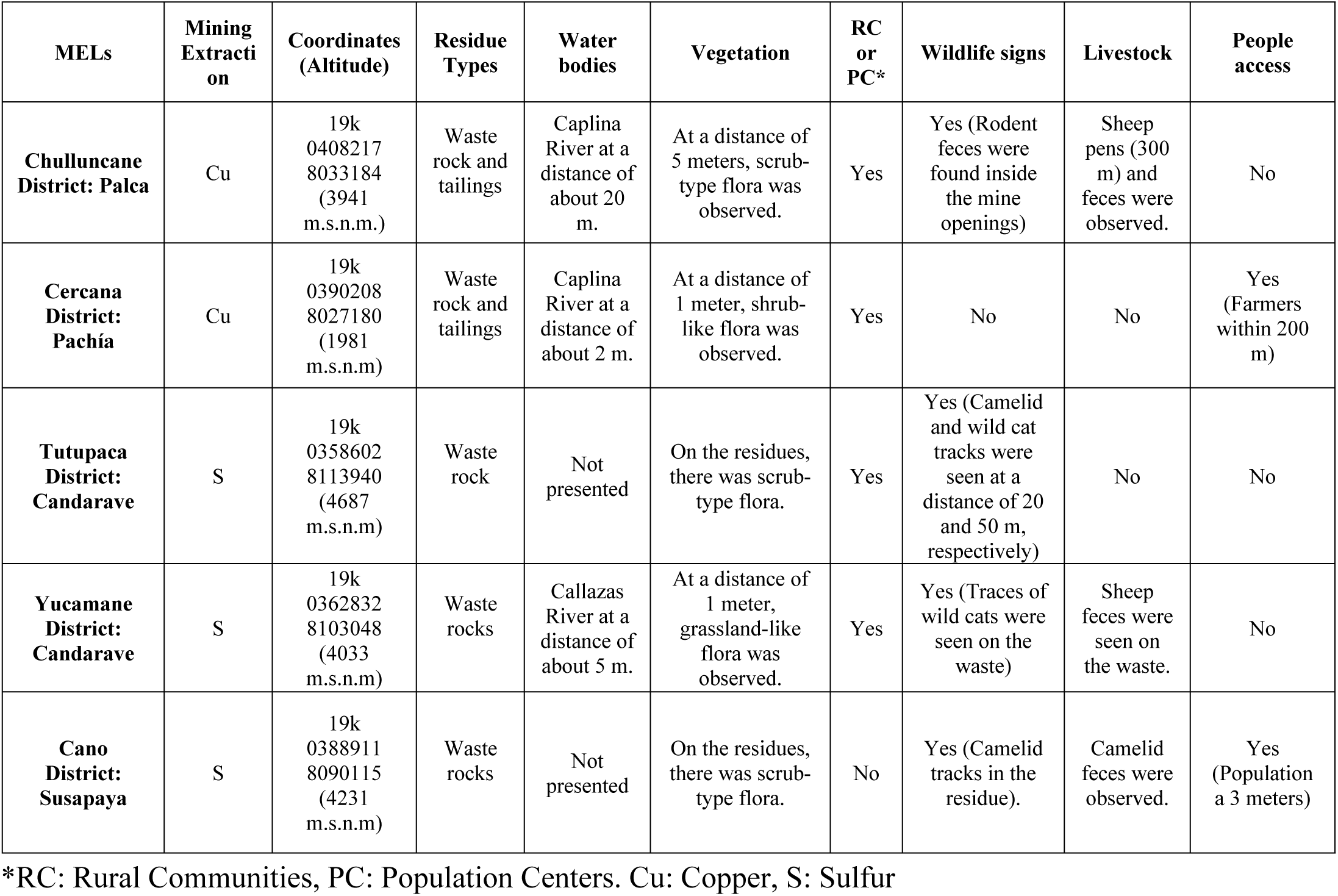
Characterization of environmental liabilities.

| MELs | Mining Extraction | Coordinates (Altitude) | Residue Types | Water bodies | Vegetation | RC or PC* | Wildlife signs | Livestock | People access |
| --- | --- | --- | --- | --- | --- | --- | --- | --- | --- |
| Chulluncane District: Palca | Cu | 19k 0408217 8033184 (3941 m.s.n.m.) | Waste rock and tailings | Caplina River at a distance of about 20 m. | At a distance of 5 meters, scrub-type flora was observed. | Yes | Yes (Rodent feces were found inside the mine openings) | Sheep pens (300 m) and feces were observed. | No |
| Cercana District: Pachía | Cu | 19k 0390208 8027180 (1981 m.s.n.m) | Waste rock and tailings | Caplina River at a distance of about 2 m. | At a distance of 1 meter, shrub-like flora was observed. | Yes | No | No | Yes (Farmers within 200 m) |
| Tutupaca District: Candarave | S | 19k 0358602 8113940 (4687 m.s.n.m) | Waste rock | Not presented | On the residues, there was scrub-type flora. | Yes | Yes (Camelid and wild cat tracks were seen at a distance of 20 and 50 m, respectively) | No | No |
| Yucamane District: Candarave | S | 19k 0362832 8103048 (4033 m.s.n.m) | Waste rocks | Callazas River at a distance of about 5 m. | At a distance of 1 meter, grassland-like flora was observed. | Yes | Yes (Traces of wild cats were seen on the waste) | Sheep feces were seen on the waste. | No |
| Cano District: Susapaya | S | 19k 0388911 8090115 (4231 m.s.n.m) | Waste rocks | Not presented | On the residues, there was scrub-type flora. | No | Yes (Camelid tracks in the residue). | Camelid feces were observed. | Yes (Population a 3 meters) |

\*RC: Rural Communities, PC: Population Centers. Cu: Copper, S: Sulfur

A concerning aspect is the presence of important rivers near the study areas, namely Caplina, Callazas, Yungani, and the proximity of peasant communities Turunturo and Palca to the study areas (Fig 2).

**Fig 2.**
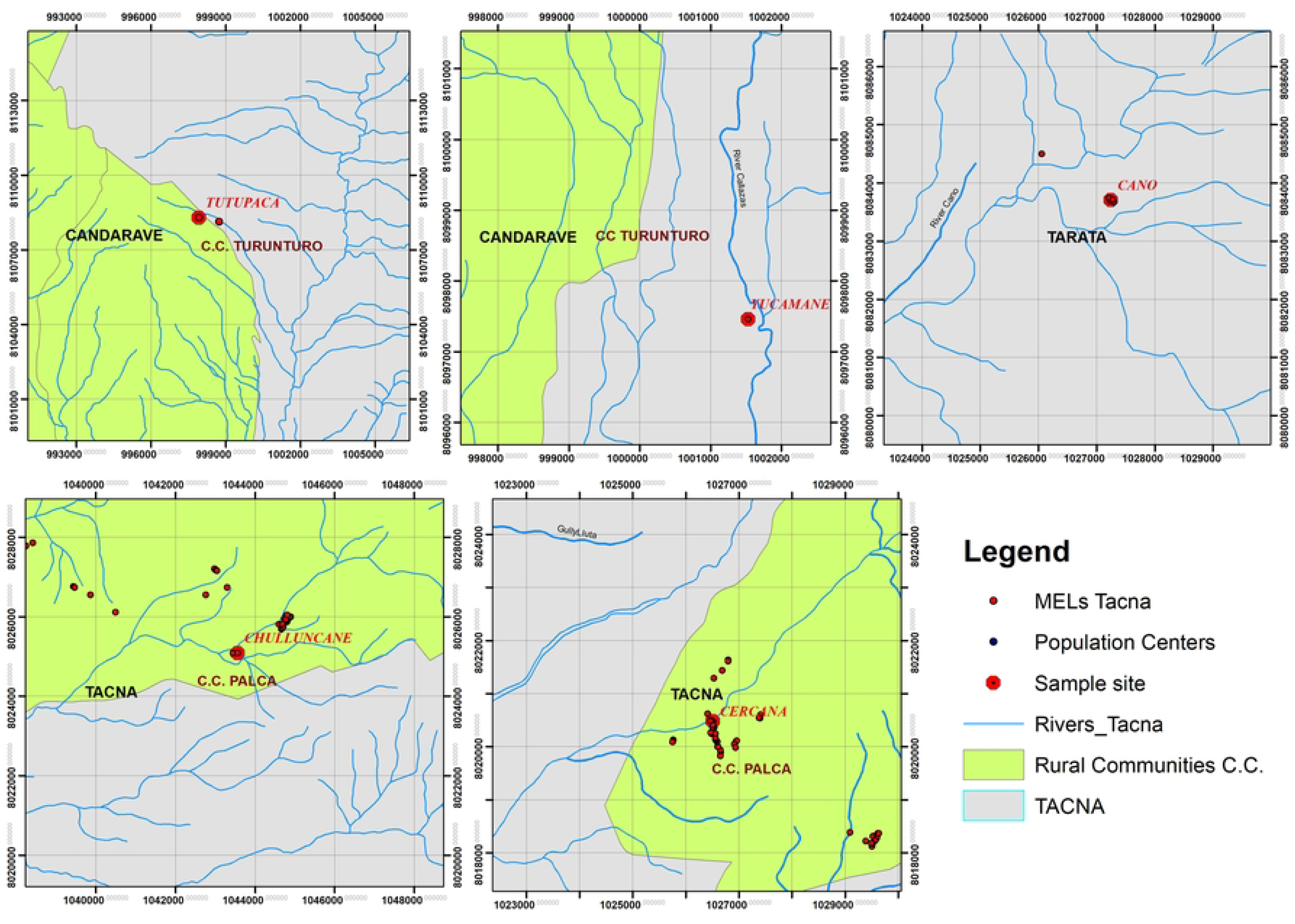
Localization of rural communities near environmental liabilities and bodies of water. Sample site: Tutupaca, Yucamane, Cano, Chulluncane, Cercana. Rural Communites: C.C. Turunturo and C.C. Palca

**Fig 3.**
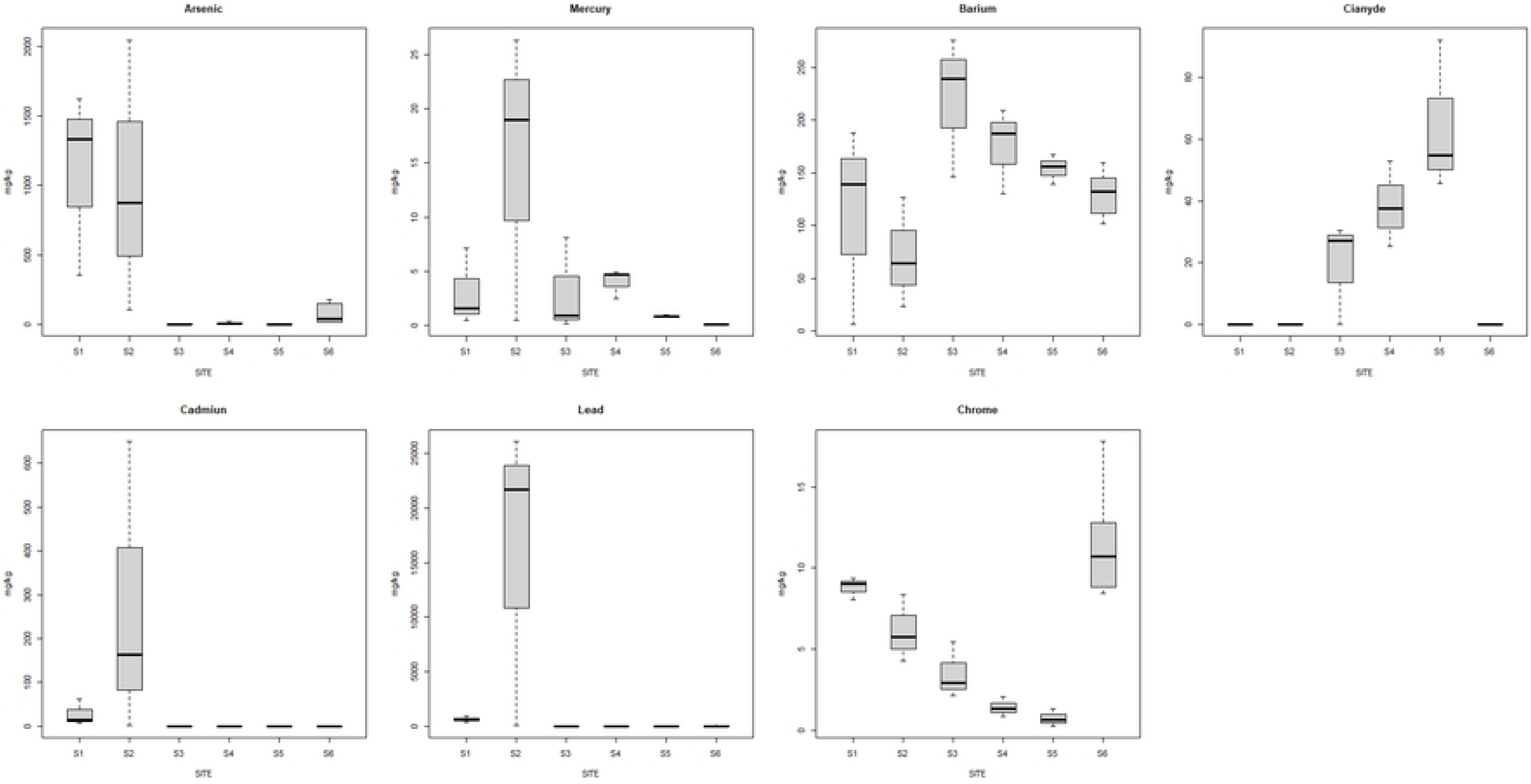
The concentration of elements analyzed for each MELs. S1: Chulluncane, S2: Cercana, S3: Tutupaca, S4; Yuca-mane, S5: Cano, S6: Reference Soil

### Soil and Potentially Toxic Elements Analysis

The average concentration of pollutants is presented in Table 2. Concentrations varied significantly depending on the sampling area, except for hexavalent chromium, which showed no differences between sites as it was below the established maximum limits. The content of various heavy metals and metalloids, as well as free cyanide in the soils, were classified in the following descending order: Chulluncane: As > Pb > Ba > Cd > Cr > Hg > CN^-^ > Cr^+6^, Cercana: Pb > As > Cd > Ba > Hg > Cr > CN^-^ > Cr^+6^, Tutupaca: Ba > CN^-^ > Pb > As > Cr > Hg > Cr^+6^ > Cd, Yucamane: Ba > CN^-^ > As > Pb > Hg > Cr > Cr^+6^ > Cd, Cano: Ba > CN^-^ > Pb > As > Hg > Cr > Cr^+6^ > Cd. It is worth noting that the concentrations of five toxic elements, including As, Pb, Cd, Barium, and free Cyanide, were considerably higher in most soil samples compared to the initial native values. Each zone has its own climatic and topographic conditions and covers an approximate area of 10 hectares. The extremely acidic pH values (<2.1) observed at Tutupaca, Yucamane, Cano, and Chulluncae are alarming. However, the pH near Cercana ranged between 4 and 5, possibly due to climatic fluctuations. Environmental liabilities with acidic pH can accelerate soil acidification processes due to anthropogenic activities and the geological characteristics of the region (71). According to Hammarström et al. (2003), in acidic conditions (pH < 4.0), pH significantly influences the adsorption of heavy metals (72). It is crucial to consider the pH factor, as acid mine drainage can cause the acidification of surface waters, which in turn may lead to the collapse of regional ecosystems and soil desertification (28).

**Table 2.**
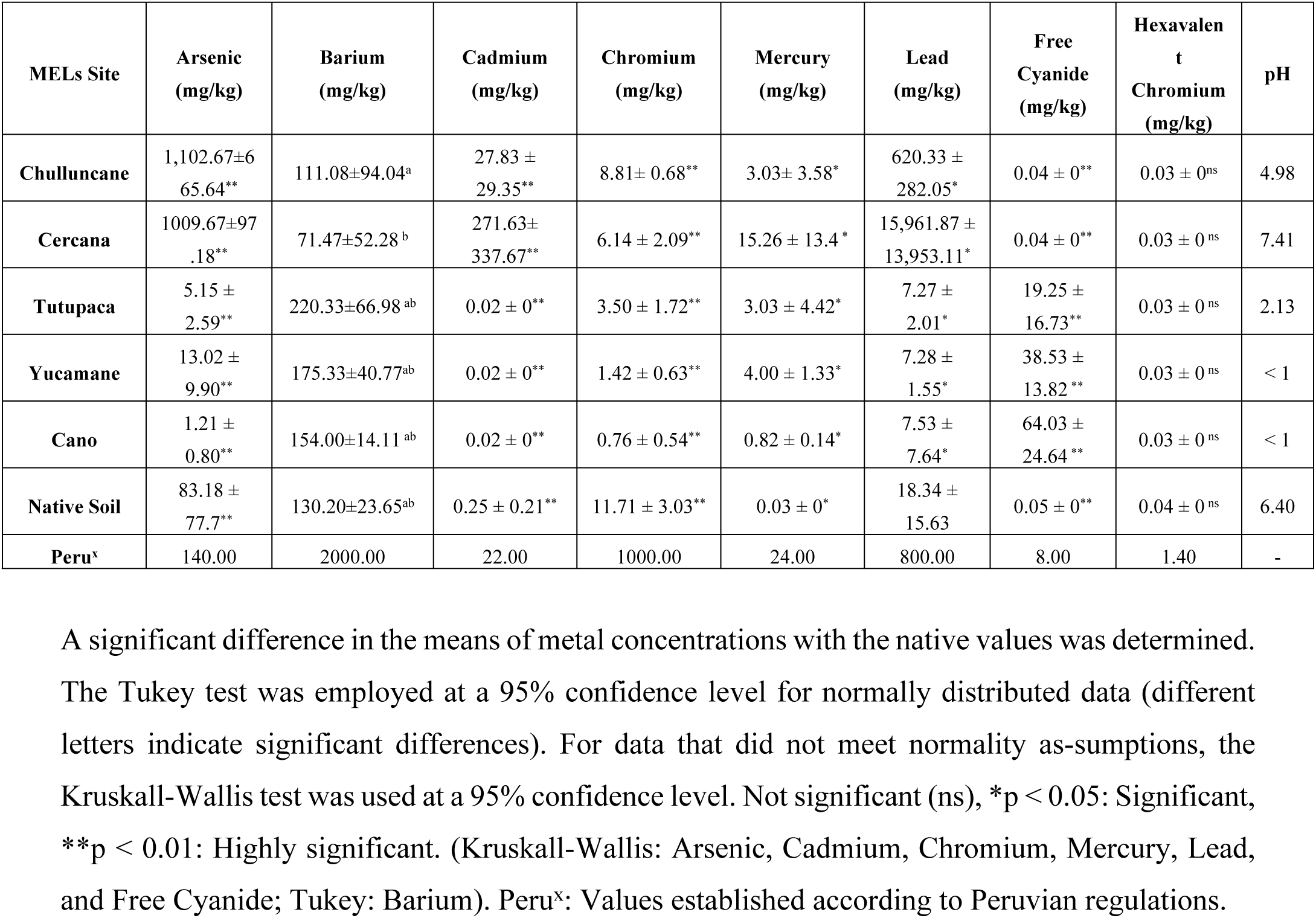
Concentration of Potentially Toxic Elements (mg/kg) in the environmental liabilities.

All values were compared against Peruvian regulations and native soil values. It was determined that the Environmental Liability (MEL) in the vicinity (MEL Cercana) has high levels of arsenic, cadmium, and lead. Meanwhile, the Environmental Liability in Chulluncane (MEL Chulluncane) exhibited elevated levels of arsenic and cadmium exceeding the established Maximum Permissible Limits (MPLs). In the Environmental Liabilities of Tutupaca, Yucamane, and Cano, there is a high concentration of free cyanide, as shown in Table 2 and Fig 2, surpassing the MPLs set by soil standards.

The toxic elements associated with mining environmental liabilities reported in this study represent a source of pollution that raises concerns within the community due to the risks they pose to the ecosystem. In Table 3, various global research endeavors are presented, detailing the sampling locations, mining extraction processes, and the studied contaminant metals, akin to our own investigation

**Table 3.** Soil contaminants from mining activities.

| Country | City | Sampling location | Mining Extraction | Pollutants | Reference |
| --- | --- | --- | --- | --- | --- |
| Peru | Tacna | Soils Contaminated by Mining Environmental Liabilities | Cu, S | As, Cd, Pb, Hg, CN<sup>-</sup> | This study |
| Perú | Cajamarca | Soils Contaminated by Mining Environmental Liabilities | - | As, Ag, Cd, Cu, Pb, and Zn | (8) |
| **China** | Daye | Actividades de fundición | Cu | Cd, Cu, Pb, As | (19) |
| **España** | Madrid | Abandoned Monica mine | Minerals (arsenopyrite, matildite and sphalerite) | As, Cu, Zn, Cd, Pb, W, Ag, Fe | (73) |
| **México** | Santa Maria de la Paz | Mining Area | Pb-Zn-Ag (Cu-Au) skarn ore system | As, Cu, Pb y Zn, | (74) |
| **Ghana** | Tarkwa, Ashanti, Atewa | Mining areas | Au | As, Cd, Hg, Zn, Co, Cu, Mn, Fe, Al, V, Cr, Pb | (75) |
| **China** | Hunan, | Nonferrous metal mine | Zn, Pb | Zn, Pb, Cd, Cu, As | (76,77) |
| **Turquía** | Kutahya | Contaminated soils by mining area | Ag | As, Ag, Pb | (78) |
| **Venezuela** | Bolívar | Soils Of The Mining Sector | Au | Hg | (79) |
| **Kosovo** | K. Mitrovica | soil of mining centres | Pb, Zn, As, Cd | As | (80) |
| **Colombia** | In northern Colombia | Soils Contaminated with Mining Activities | Au | Hg, Pb y Cd | (81) |
| **Eslovaquia** | Central Spiš | Soil Heavy Metal Pollution | Cu, Hg | Hg, Zn y Cd | (82) |
| **India** | Karnataka | Tailings | Au | Cd, Cu, Pb, Zn | (52,83) |
| **Polonia** | Zloty Stok | Mining soil | Au, As | As | (84) |
| **Perú** | Pasco | soil in mining towns | Ag, Au | Pb | (85) |
| **Kosovo** | Mitrovica | Soil and tailing | Pb, Cd, Zn | Pb | (86) |
| **Colombia** | Sabana de Bogotá | Soil | - | Pb | (87) |
| **Chipre** | Lefke | Soil and plant | Cu | Fe, Zn, Pb, Cd, Cu, Ni, Cr | (88) |
| **México** | San Francisco of Oro (Chihuahua) | Soil | - | Pb, Zn, Cd, As | (89) |
| **Etiopía** | Adola | Sediment, water and plant | Au | Al, Mn, Fe, As, Ni, Cr, Cu, Co, Pb, W, Sb, Mo, Zn, V | (90) |
| **Polonia** | Tarnobrzeg County | Mining Soil Substrate | S | N, Ca, Mg, Al, S | (91) |
| **China** | Hainan | Soils, mine tailing, and groundwater samples | Au | Pb, As, Cd, Hg, CN- | (92) |

#### Calculation of Environmental Indices

To assess pollution in each area, different pollution indices were calculated. These potential pollution indices were computed using the concentration of an element in the topsoil layer relative to the concentration of a reference element.

#### Geoaccumulation Index

Calculations for the Geoaccumulation Index reveal high levels of contamination in certain Environmental Liability sites (Table 4). Specifically, the Chulluncane MEL shows extreme contamination levels for Cd and Hg (5.235 and 5.229, respectively), and strong contamination for As and Pb (3.807 and 3.122, respectively). The Cercana MEL displays extreme contamination levels for Hg, Pb, and Cd (7.011, 6.585, and 5.713, respectively), along with high to extreme concentrations of As (4.301).

**Table 4.** The study includes the Geoaccumulation Index of environmental liabilities.

|  | Geoaccumulation Index |  |  |  |  |
| --- | --- | --- | --- | --- | --- |
| Pollutant | Chulluncane | Cercana | Tutupaca | Yucamane | Cano |
| As | 3.807 | 4.301 | -2.583 | -4.632 | -7.777 |
| Ba | -2.135 | -1.414 | 0.341 | -0.339 | -0.367 |
| Cd | **5.235** | **5.713** | -2.392 | -2.755 | -3.832 |
| Cr | -1.603 | -1.701 | -2.028 | -3.601 | -4.348 |
| Hg | **5.2289** | **7.011** | 4.445 | **6.411** | 4.174 |
| Pb | 3.122 | **6.585** | -0.682 | -1.055 | -1.191 |
| $\text{CN}^-$ | -0.907 | -0.907 | **5.421** | 8.94 | **9.671** |
| $\text{Cr}^{+6}$ | -1 | -1 | -1 | -1 | -1 |

$\leq 0$ , Unpolluted; $0 \leq I\text{-geo} \leq 1$ , Unpolluted to moderately polluted; $1 \leq I\text{-geo} \leq 2$ , Moderately polluted; $2 < I\text{-geo} \leq 3$ , Moderately to strongly polluted; $3 \leq I\text{-geo} \leq 4$ , Strongly polluted; $4 \leq I\text{-geo} \leq 5$ , Strongly to extremely polluted; and $5 < I\text{-geo}$ , Extremely high polluted.

In Tutupaca, contamination is extreme only for free CN^-^ (5.421) and high to extreme for Hg (4.445). The Yucamane MEL exhibits extreme contamination levels for free CN^-^ and Hg (8.94 and 6.411, respectively), while the Cano MEL shows extreme contamination for free CN^-^ (9.671) and high to extreme contamination for Hg (4.174).

#### Contamination Factor

Using the Contamination Factor (*C_f_*) index, it is necessary to quantify the contamination by hazardous compounds, as illustrated in Table 5.

**Table 5.**
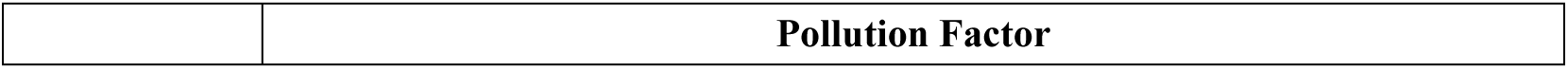
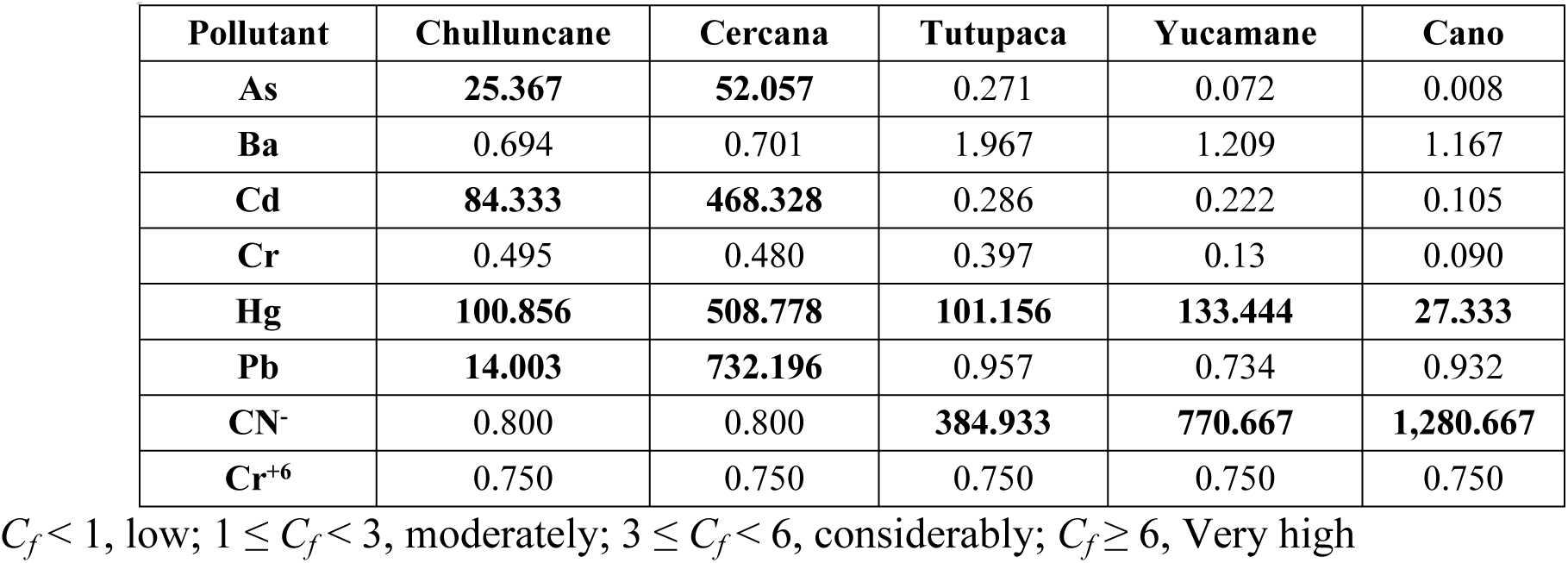
Contamination Factor Index of Environmental Liabilities Included in the Study.

As observed in the Environmental Liabilities (MELs), contaminants exceeding the established *C_f_* value of ≥ 6 resulted in a high contamination index. In Chulluncane, we have Hg > Cd > As > Pb. In Cercana, Pb > Hg > Cd > As. In Tutupaca, CN^-^ > Hg. In Yucamane, CN^-^ > Hg. Finally, in the MEL located in Cano, CN^-^ > Hg. The prevalent contaminants in each environmental liability are Hg, Pb, and CN^-^, making them among the most concerning.

The higher *C_f_*values indicate a significant role played by anthropogenic contaminants, while lower *C_f_* values highlight the geological distribution of components in the soil.

#### Pollution Load Index

The Pollution Load Index indicates the degree of contamination in each evaluated area (Fig 4). The descending order of contamination levels is as follows: Cercana (12.3) > Chulluncane (4.3) > Yucamane (1.8) > Tutupaca (1.95) > Cano (1.02), with an average of 4.3. All assessed sites have values greater than 1, indicating contamination of the Environmental Liabilities (MELs). The contamination index is based on the concentration of metals and the total number of analyzed pollutant parameters found in a sample, serving as an indicator of severe contamination due to the presence of toxic compounds. These compounds may originate from mineral extraction processes, copper (Chulluncane and Cercana), and sulfur (Tutupaca, Cano, and Yucamane), and their presence can lead to increased pollution.

**Fig 4.**
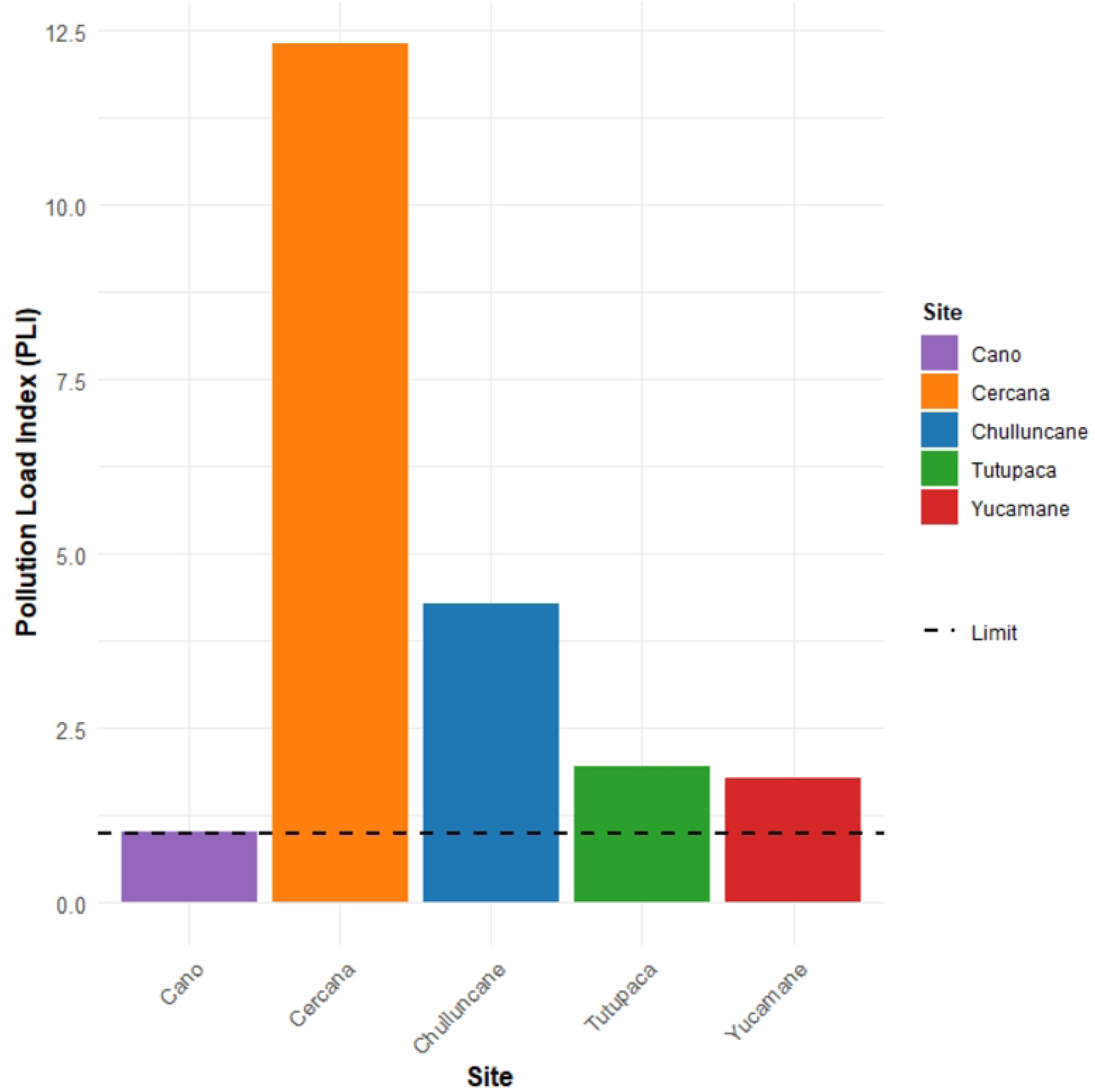
Pollutant Load Index (*PLI*) for five sites. The mean *PLI* values indicate that four sites surpass the pollution threshold of 1, with **Cercana** showing a notably high mean *PLI* of **12.3**, indicating severe contamination. **Chulluncane** follows with a mean *PLI* of 4.27, while **Tutupaca** (1.96) and **Yucamane** (1.79) exhibit moderate pollution. **Cano** is slightly above the limit with a mean *PLI* of 1.01.

#### Degree of contamination, modified degree of contamination, and potential ecological risk

The degree of contamination (*Cdeg*) and modified degree of contamination (*mCdeg*) indicated, in decreasing order, contamination values for the study zones as follows: Cercana (2,640 and 330) > Cano (1,311 and 163) > Yucamane (907 and 113) > Tutupaca (490 and 68) > Chulluncane (227 and 28) (Fig 5). In all cases, *Cdeg* and *mCdeg* values exceed the established thresholds of 24 and 36, respectively, indicating a high degree of contamination, mainly due to anthropogenic pollution generated by As, Ba, Cd, Hg, Pb, and CN^-^.

**Fig 5.**
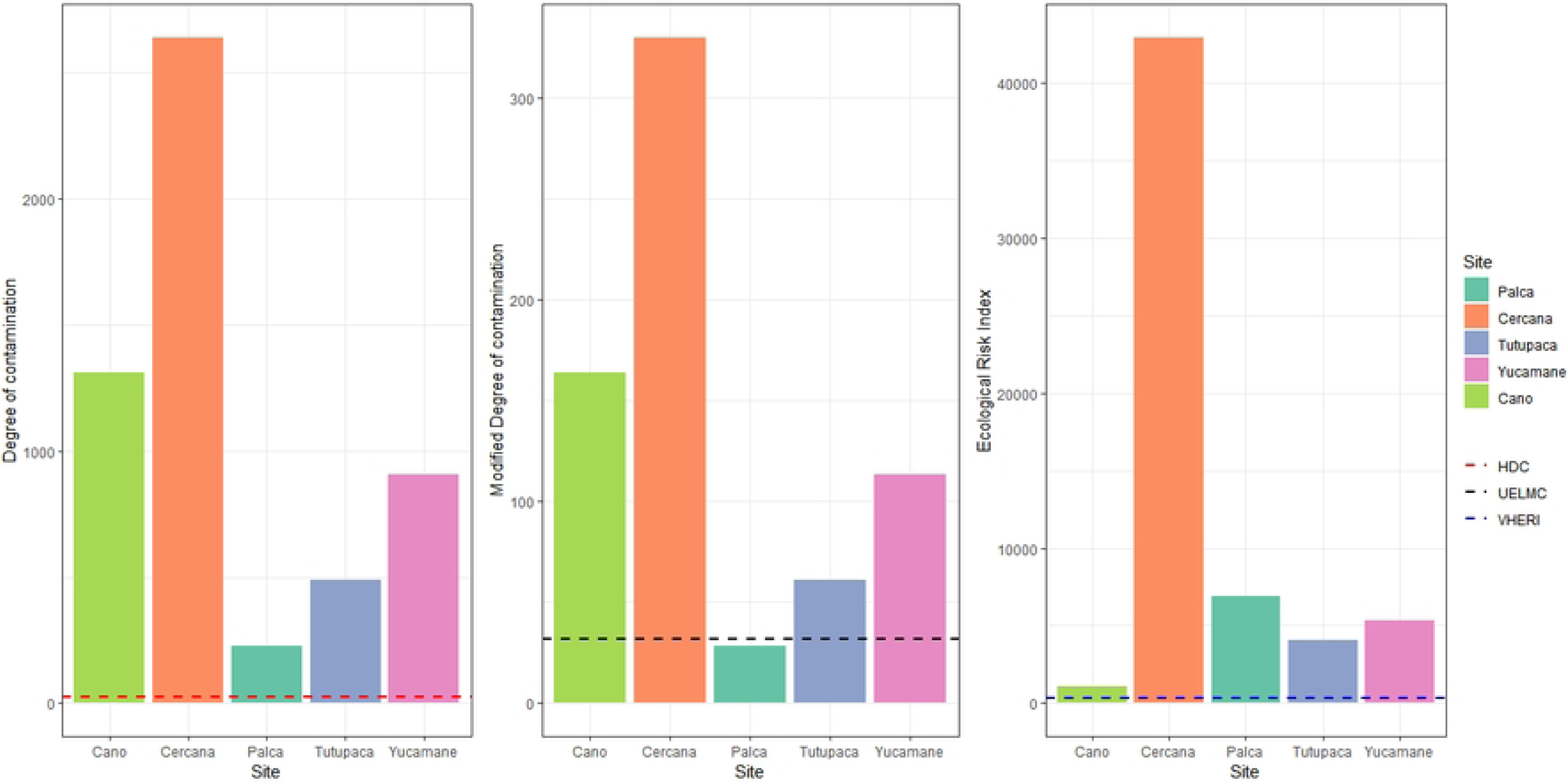
Potential ecological risk. The figure illustrates the Degree of Contamination (*Cdeg*), Modified Degree of Contamination (*mCdeg*), and Ecological Risk Index (*ERI*) across five sites: Cano, Cercana, Chulluncane, Tutupaca, and Yucamane. The dotted lines mark the thresholds for High Degree of Contamination (*Cdeg* > 24), Ultra-high Degree of Contamination (*mCdeg* ≥ 32), and Very High Ecological Risk (*ERI* > 600). Cercana exhibits exceptionally high values in all metrics, exceeding these thresholds, indicating severe contamination and ecological risk. Cano, Chulluncane, and Yucamane show moderate levels of contamination, while Tutupaca remains below most critical thresholds.

Regarding potential ecological risk (Fig 5), the presented values exceed 600, indicating a very high potential ecological risk index or a severe ecological contamination level, due to contamination produced by As, Cd, Cr, Hg, and Pb.

### PCA of Metals and Site Distributions

The first two axes of the PCA represented 76.6% of the total variance (PC1 explained 52.9%, and PC2 explained 23.7%). The first axis clearly separated the three different sites in the PC space, which indicated that they showed statistical differences among soils S1, S2, S3, S4, and S5. In detail, sites S1 and S2 are positively and negatively distributed along PC1, respectively, while S3, S4, and S5 are mainly distributed negatively along PC2 (Fig 6).

**Fig 6.**
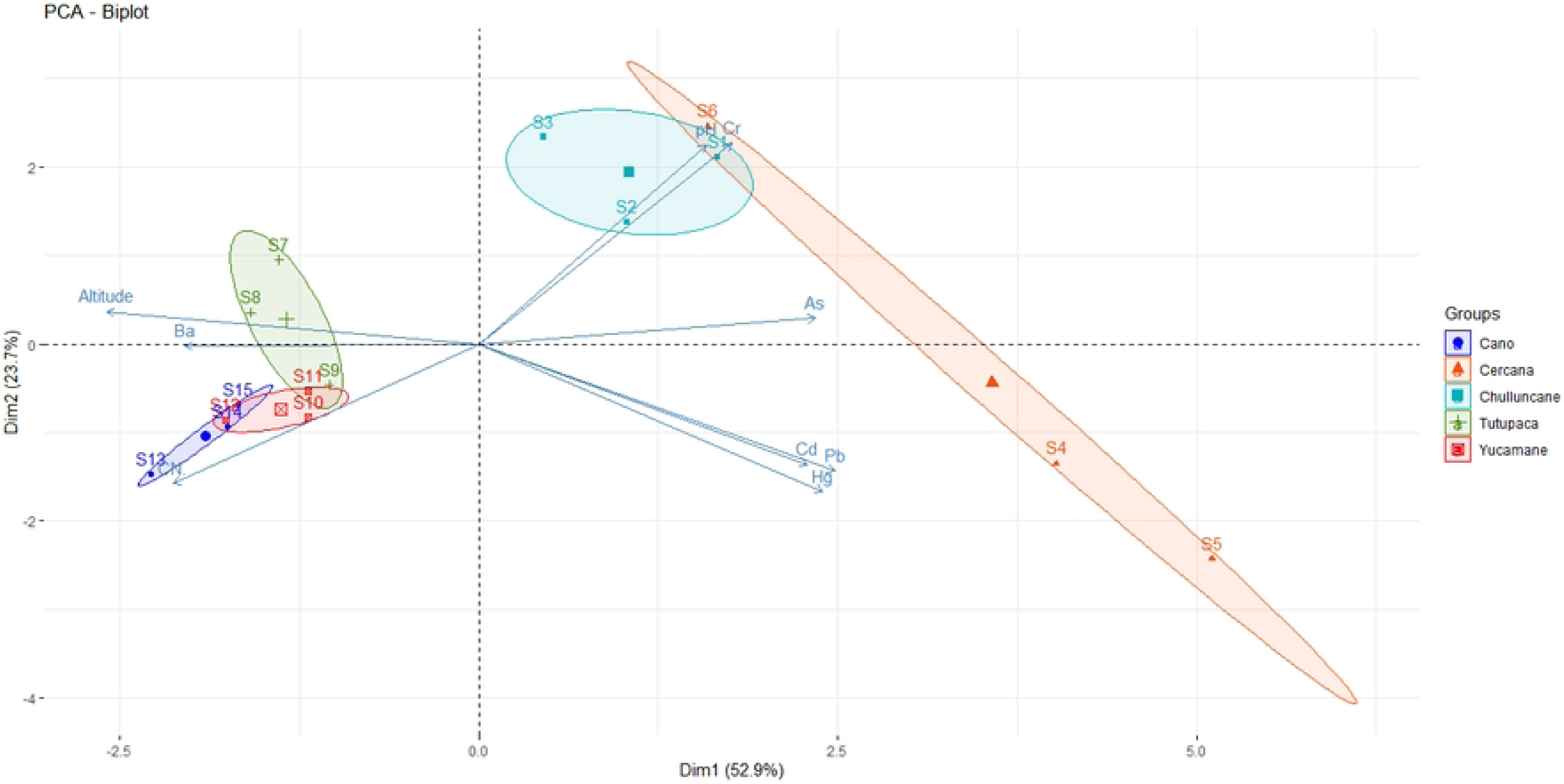
Principal Component Analysis (PCA) graphs of metal concentrations (As, Cd, Cr, Pb, Br, Hg, and CN^-^) and their spatial distribution in environmental sites of S1: Cercana, S2: Chulluncane, S3: Tutupaca, S4: Yucamane, S5: Cano. Circular lines on the PCA graphs overlap to indicate sampling sites in the same area.

### Correlation among Parameters

A Spearman correlation analysis was conducted among potentially toxic elements to investigate the relationship of metals in environmental liabilities (Fig 7). Strong positive correlations were observed between Cd/Hg (r=0.87), Cd/Pb=(r=0.78), Hg/Pb (r = 0.91), CN^-^/Cr (r = -0.77), As/Cr (r=0.59, As/Hg (r=0.55) and CN^-^/pH (r=0.71), indicating a possible co-occurrence of these heavy metals, suggesting a common source or similar environmental processes. Chromium (Cr) shows a positive correlation with pH (0.79), which could affect its mobility in the soil. On the other hand, significant negative correlations were identified, such as Cr with CN (-0.77), Pb (-0.76), and Hg (-0.67) with altitude, suggesting that these metals decrease in concentration at higher elevations. These correlations highlight the importance of geographic and chemical factors in the distribution of contaminants, providing a crucial basis for environmental management in affected mining areas.

**Fig 7.**
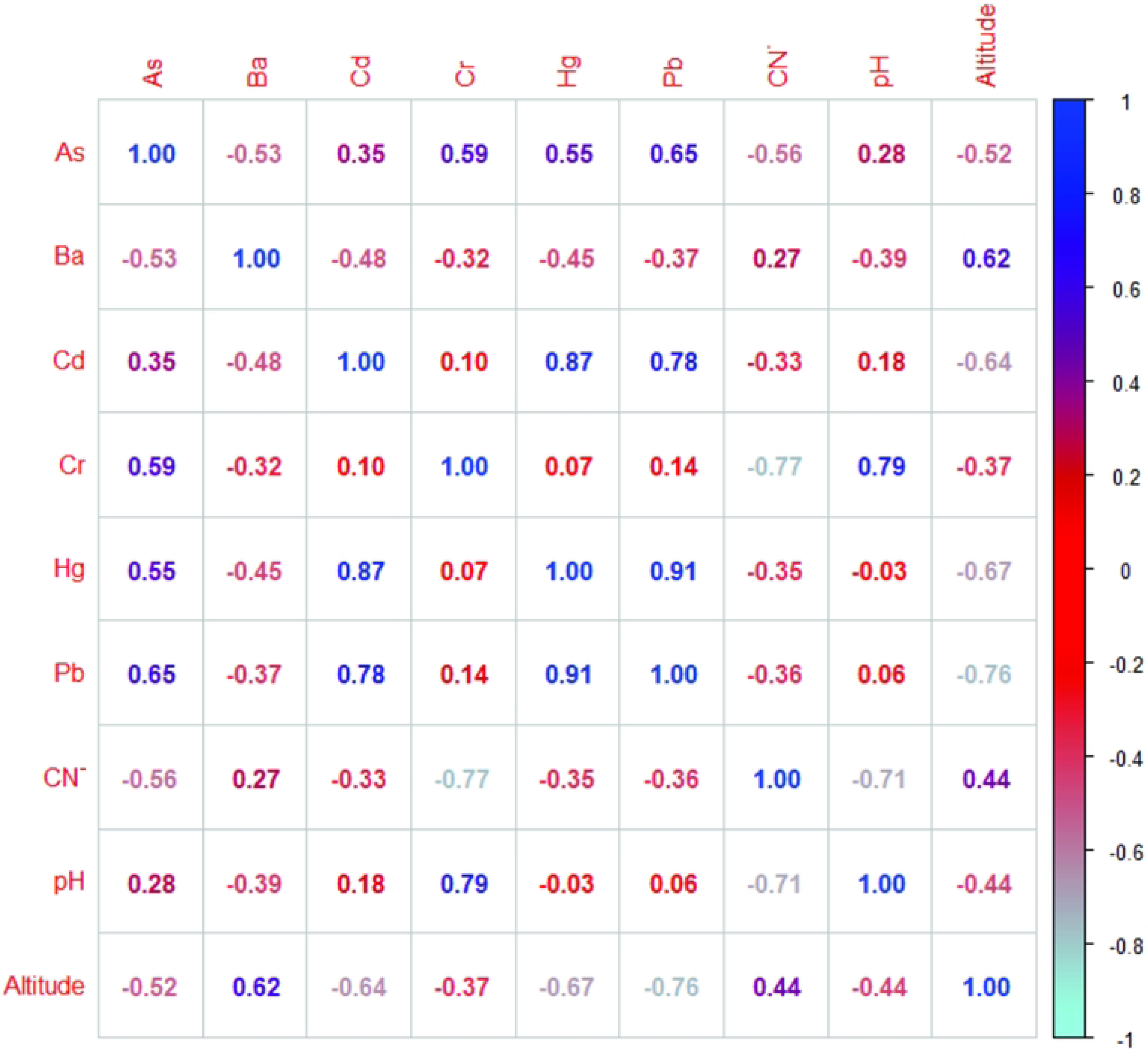
Spearman rank order correlation for selected parameters analyzed in soil samples from mining environmental liabilities.

## Discussion

The environmental liabilities in this study represent an underlying issue as they contain significant quantities of potentially toxic elements that can alter the ecosystem and give rise to social conflicts, particularly ^-^due to their proximity to rural communities. In some locations, agricultural and livestock activities were recorded, along with the presence of rivers. The environmental liabilities (MELs) analyzed in this study are heavily contaminated with Pb, As, Cd, and CN-. These residues could have originated during mining exploitation processes (35), potentially extending over time to various ecosystems. In another study conducted on potentially toxic elements in environmental liabilities in the Peruvian Andes, concentrations of six elements were found, Pb, Zn, As, Cu, Ag, and Cd (1); these elements are similar to those reported in our investigation. Additionally, mining tailings revealed concentrations of arsenic (As), cadmium (Cd), chromium (Cr), cobalt (Co), copper (Cu), lead (Pb), manganese (Mn), nickel (Ni), and zinc (Zn) in soil samples, varying with pollution factors (4). Furthermore, ecosystems in abandoned mining sites were reported to be affected by acid mine drainage, with the presence of metals such as Al, Ca, Co, Cu, Fe, Mg, Mn, Ni, and Zn (39). Studies on waste disposal sites also found contamination by Pb, Zn, and Cu, which were the most concerning and pose significant environmental threats (58).

The averages of pollutant elements were found to be considerably higher at most of the assessed sites compared to reference values. Elevated concentrations of arsenic and cadmium were observed at the Chulluncane and nearby mines, lead at the nearby mine, and free cyanide at the Tutupaca, Yucamane, and Cano mines, surpassing Peruvian regulatory limits. This could be attributed to the processes involved in the extraction of copper and sulfur minerals, which undergo milling, and smelting, among others. It is important to emphasize that each process generates some form of waste (9). Additionally, industrial waste, landfills, extraction waste materials, vehicle emissions, waste incineration, construction, demolition, incomplete mine closure, and inadequate mineral recovery, coupled with operators’ non-compliance with environmental policies (93), serve as sources of contamination.

The input of these contaminants into water bodies (11), soil, and cultivated lands (22) must be considered in the evaluated MELs, as it may lead to alarming damage to the ecosystem (72,94). Biodiversity represents a threat to health through ingestion, inhalation, and contact (95–98), creating carcinogenic risks to the lungs, liver, kidneys, bladder, and breasts, as well as cardio-pulmonary effects and alterations in gene expression (21,97,99–103). Furthermore, there are associated risks of Alzheimer, osteoporosis, diarrhea, vomiting (104–109), and impacts on the social well-being of communities (28,33,89,104,110).

High concentrations of toxic metals and metalloids, such as As, can lead to adverse health effects, including a potential carcinogenic risk (111). As is associated with copper and gold-copper minerals (112). It has also been reported that arsenic concentrations are elevated in abandoned mines and areas with tailings reservoirs (74,113). Additionally, a mining site in the province of Salamanca, Spain, exhibited extreme As levels (> 1,000 mg/kg), possibly attributed to the presence of sulfide minerals in altered granites or metamorphic rocks (74,113).

On the other hand, Razo et al. (2004) reported As concentrations of 29,000 mg/kg and 28,600 mg/kg in areas of former mining, where tailings reservoirs come into contact with air and soil. A study conducted in Peru, specifically in the Cajamarca region, found elevated concentrations of Pb and As exceeding Peruvian standards, while Cd levels remained within permissible limits (8). However, Piura, a mine dedicated to copper ex-traction reported high levels of Cd, As, and Pb above permissible limits (114). Our findings indicate that in Tacna, Cd levels reached up to 650 mg/kg, As reached 2,046 mg/kg, and Pb reached 16,000 mg/kg. Furthermore, concentration levels are much higher, including Cd (650 mg/kg), free cyanide (92 mg/kg), and Pb (2,6131 mg/kg). In the Cajamarca mine, Pb levels also reached up to 16,000 mg/kg (114), exceeding values reported in other studies on heavy metal contamination in mining environmental legacies, such as Carolina – Cajamarca, and the Cabeza Rajao mine in Spain, with Cd (41 mg/kg), As (315 mg/kg), and Pb (8 mg/kg) values (26).

On the other hand, elevated concentrations of Pb and Cd may result from a significant amount of mining waste slag or the mining extraction process itself, akin to findings reported in other studies (7,83,92,115). Furthermore, the high availability of Hg in the environment is associated with sulfur concentrations and pH, leading to increased bioavailability, potentially forming soluble or solid complexes (79).

Additionally, pollution indices such as *I-geo, C_f_, PLI, Cdeg, mCdeg*, and the ecological risk index allowed for a comparison of contamination levels in an area with reference values. All conducted analyses indicate that the zones exhibited some level of contamination caused by one or more toxic elements, linked to availability and extractive activity. An aggravating factor in this situation is the acidic pH of the study areas, indicating a deterioration of MELs (Metallic Elements Limits). The effect of rain runoff and nearby rivers can mobilize pollution towards the surrounding communities, such as the rural communities of Turunturo and Palca, posing a potential risk of infiltration of mining residues and interaction with natural water courses (116). It is also crucial to consider the possibility of metal mobilization due to biogeochemical processes associated with them, coming into contact with water bodies and soils (117,118).

Specifically, the values PLI (Fig 4), *Cdeg, mCdeg* (Fig 5), indicates a marked contamination in all study areas. The *PI* values for As, Cd, Pb, and CN^-^ were >6, indicating high contamination of these pollutants in the MELs. Additionally, it allowed the inclusion of Hg as a contaminant (*PI* > 6), indicating a very high contamination factor. The *PLI* indicates that all environments deteriorate, and *Cdeg, mCdeg* indicates a high degree of contamination. Therefore, these pollutants may have adverse effects on health. This implies that these contaminants were higher than in reference areas, suggesting that these elements originated from anthropogenic activities during the time the mine was operational. This suggests the need for monitoring (119) to verify the presence of these PTEs in nearby soils and the sediment of the water network (35).

To assess the potential impact of PTEs on biota, *RI* and *Eri* were used, complementing information from *PI, PLI, Cdeg,* and *mCdeg*. *IR* values > 600 indicated a very high ecological risk or severe ecological contamination, representing an ecological risk to the biota of this location and a considerable ecological risk to the surrounding area. These findings suggest that contamination may result from anthropogenic activities such as industrial mining, accumulation, and discharge of mining waste, as reported by other authors (77,83,92,120,121).

Several studies have indicated ecological risks ranging from moderate to high due to the presence of potentially toxic elements (4,29,38,39,58,122,123). Additionally, our research has provided a preliminary assessment of the risk associated with the presence of mining environmental liabilities, which signals a high level of pollution in proximity to both populations and bodies of water. This risk could be mitigated if these liabilities were situated away from populated areas and bodies of water (29). Other studies conducted on abandoned gold mine tailings pose severe ecological and human health risks linked to exposure to heavy metals and metalloids released into the environment.

High concentrations of As, Cd, Co, and Ni in the soil may pose unacceptable risks to human populations (4), affecting ecosystems impacted by acid mine drainage and metal-enriched conditions (Al, Ca, Co, Cu, Fe, Mg, Mn, Ni, and Zn) found in adjacent tailings in semi-humid climatic conditions (39). The primary concern lies in contamination by Pb, Cu, and Cr due to human activities (124).

In the present study, pollutant concentrations in mining environmental residuals and PCA results demonstrated distinct sources of contamination. Nevertheless, sites S1, S2, S3, S4, and S5 exhibited pollution attributed to anthropogenic activities, suggesting that these metals may have originated from common sources (125–128). Some research also indicates that they may derive from parental materials through lithogenic and pedo-genic processes (129).

Furthermore, it is noteworthy that the primary sources of Cu, Mn, and Zn contamination also appear to be linked to human activities, such as agriculture, urbanization, and industrialization, significantly impacting the investigated areas.

The observed positive correlation between Cd/Hg (r=0.87), Cd/Pb=(r=0.78), Hg/Pb (r = 0.91), As/Cr (r=0.59, As/Hg (r=0.55) and CN-/pH (r=0.71). This suggests a possible co-occurrence of heavy metals, which could indicate a common source or the influence of similar environmental processes (125–128). Additionally, the Spearman correlation value with the studied metals indicates that the transport of contaminants (toxic metals) is related to the evaluated variable (130). Similar results were observed at different sites, where correlations between metal concentrations are influenced by geoclimatic behavior. Furthermore, potentially toxic elements have been found to persist in the environment for extended periods without significant biodegradation following their introduction from anthropogenic sources. This poses a substantial risk to residents near environmental liabilities and landfills, potentially impacting the health of the surrounding ecosystem.

In this regard, it is essential to comprehend the distribution of toxic compounds in environmental liabilities to guide decision-makers in establishing effective environmental protection measures. In our study, extremely high concentrations were observed for Pb, As, Cd, Hg, and CN^-^. According to ecological risk indices, the order for implementing mitigation measures or a recovery management plan in this area would be as follows: Cercana, Chulluncane, Yucamane, Tutupaca, and Cano. These sites need proper remediation to prevent them from becoming sources of contamination for the community through polluted soils and sediments, posing risks to both ecology and human health of the inhabitants (35).

The findings in this study reveal high to elevated levels of contamination, suggesting that soils associated with mining environmental liabilities are unsuitable for agricultural use in the investigated areas (131). Due to the high toxicity, it is imperative to reinforce measures to ensure compliance with environmental regulations and promote scientific research in this field (35). Additionally, implementing a monitoring program is crucial to reduce pollution in the environment, as all these contaminants can lead to various health problems (132).

Therefore, it is crucial to consider risk assessment in ecosystems and the characterization of MELs in order to formulate approaches, ecological techniques, and scenarios for crafting policies that promote their economically viable mitigation, containment, and elimination (133). For the treatment of potentially toxic elements (134), it is necessary to implement technologies based on bioremediation, nanotechnology, bioventilation, bioaerosolization, biostimulation, bioaugmentation, and phytoremediation, which could prove effective in mitigating pollution in the studied MELs (134–139). Additionally, alternatives like tailings pond coverages (139), soil replacement, enhancement of surface layers, compatible covers (surface coating), soil washing, the use of organic matter, and the planting of native or exotic species, among others (45,81,140–143), can seamlessly integrate with the aforementioned techniques and expedite the remediation of contaminated soils.

The resolution of this issue is intricate and multifaceted, necessitating the implementation of various technological strategies to develop short, medium, and long-term solutions, enabling the ecosystem to recover its ecological integrity sustainably and scientifically soundly (144).

All the studied liabilities lack direct or indirect responsible parties, thus the Peruvian government must take responsibility for the control and remediation of these Environmental Liabilities. However, there are no actions on the part of decision-makers, as the number of environmental liabilities is substantial, and there is no situational diagnosis guiding the government on measures to adopt in order to minimize the impacts on the environment.

## Conclusions

The quantification of pollution in the study area revealed extremely high concentrations of As, Cd, Pb, and CN^-^. Analysis of various pollution indices exposed critical conditions across all evaluated areas and identified Hg as a potentially toxic agent. These findings underscore the need to employ environmental pollution index models to accurately assess the pollution levels resulting from environmental liabilities within the ecosystem. The elevated pollution levels observed in the *PLI, Cdeg*, and *mCdeg* indices reflect severe environmental impacts.

The assessment of ecological risk indices (*IR*) and ecological risk index (*Eri*) suggests an extremely high ecological risk, indicating a severe level of contamination that poses a significant threat to local biota and a considerable risk to surrounding areas. In this context, it is crucial to establish a comprehensive monitoring program to reduce pollutant loads and identify critical contamination points to mitigate future environmental impacts. Implementing policies that promote economically viable mitigation, containment, and removal of contaminants is essential. The results of this study also provide a preliminary foundation for addressing the risks associated with mining environmental liabilities in the soils of the Tacna region. It is important to recognize that resolving this complex issue requires the adoption of short-, medium-, and long-term technological strategies to sustainably restore the ecological integrity of the ecosystem, supported by robust scientific evidence.

## Acknowledgments

This research was funded by the National University Jorge Basadre Grohmann through the ’Mining Canon and Mining Royalties Fund,’ as sanctioned by Rectoral Resolution No. 4723-2015-UN/JBG. The funding supported the project titled ’Genomic Analysis of Cyanide-Degrading Microorganisms for the Remediation of Mining Environmental Liabilities in the Tacna Region’.

## Author Contributions

**Conceptualization:** César J. Cáceda, Milena Carpio

**Data curation:** César J. Cáceda, Milena Carpio, Gisela Maraza, Gabriela de L. Fora

**Formal analysis:** César J. Cáceda, Milena Carpio

**Funding acquisition:** César J. Cáceda

**Investigation:** César J. Cáceda, Milena Carpio

**Methodology:** César J. Cáceda, Milena Carpio, Gisela Maraza, Gabriela de L. Fora, Gabriela de L. Fora, Edwin Obando, Diana Galeska Farfan

**Project administration:** César J. Cáceda, Milena Carpio

**Software:** Milena Carpio

**Supervision:** César J. Cáceda, Milena Carpio

**Validation:** César J. Cáceda, Milena Carpio

**Writing – original draft:** César J. Cáceda, Milena Carpio, Gisela Maraza, Gabriela de L. Fora, Edwin Obando, Diana Galeska Farfan

**Writing – review & editing:** César J. Cáceda, Fulvia Chiampo, Milena Carpio

